# Spatialsnake: a flexible workflow for spatial transcriptomics analysis across platforms

**DOI:** 10.64898/2026.10.27.762082

**Authors:** Zhenghao Lin, Zhihao Zhang, Nan Huang, Zixiao Lu, Siwen Xu

## Abstract

**Background:** Spatial transcriptomics technologies measure gene expression at different spatial resolutions and generate diverse data outputs. Existing workflows support a range of platforms and analyses, but combining methods and extending analyses to selected populations or tissue regions can require additional scripts and repeated data preparation.

**Findings:** We developed Spatialsnake, a flexible workflow built on SpatialData and Snakemake for analysing processed outputs from Visium, Visium HD, Stereo-seq, Xenium and MERFISH. A common command-line interface supports single- and multi-sample analysis, with adjustable parameters and saved intermediate results. Users can select populations, tissue regions or distance-based neighborhoods, repeat clustering and annotation, and map refined labels back to the full dataset. Additional modules support cell-type composition estimation, spatial-domain identification, cell– cell communication inference, regulon analysis and sample-level comparisons. We demonstrate these functions using mouse-brain Visium and colorectal cancer Visium HD datasets. Joint analysis of three colorectal cancer sections retained 1,527,987 bins and enabled further analysis of 223,632 epithelial bins, linking epithelial subpopulation annotations to their locations within the complete tissue.

**Conclusions:** Spatialsnake provides a unified workflow for spatial transcriptomics analysis across sequencing- and imaging-based platforms. Its flexible analysis modules support both whole-tissue analysis and further investigation of selected cell populations and tissue regions.

## Background

Spatial transcriptomics links gene expression to tissue location, allowing researchers to investigate how cellular composition and molecular states are organized within tissues. This spatial view can be obtained using sequencing- or imaging-based technologies, which differ in molecular coverage and spatial resolution[1]. Depending on the technology, expression is measured in spots, bins or segmented cells and is accompanied by tissue images, transcript coordinates or cell boundaries. These differences in measurement and output mean that applying common analysis methods requires different data preparation steps across platforms.

Several workflows have been developed to simplify spatial transcriptomics data processing and analysis. These workflows differ in where they begin and the technologies they support. Spacemake connects raw sequencing data processing with spatial analysis and reporting [2], whereas Panpipes supports spatial preprocessing, clustering and deconvolution alongside its single-cell multiomic workflows [3]. For image-based spatial omics, Sopa combines segmentation, aggregation, annotation and spatial analysis [4]. More specialized workflows address particular processing steps, including stPipe for generating spatial count data from sequencing reads [5] and ENACT for segmentation, bin-to-cell assignment and annotation of Visium HD data [6]. Together, these tools support a range of spatial technologies and analytical tasks, although researchers may still need to combine tools when their analyses extend beyond the coverage of a particular workflow.

The need to combine tools becomes particularly apparent when an analysis moves from tissue-wide annotation to the study of specific populations or regions. For example, researchers may wish to resolve subgroups within an epithelial population or examine the cells surrounding a tumor region. Such investigations involve selecting relevant observations, adjusting analysis settings and carrying the resulting subsets into further analyses, such as cell–cell communication inference or gene regulatory network analysis. Although methods are available for these tasks, using them together requires compatible inputs and consistent handling of sample identities, annotations and spatial coordinates. A workflow that connects these steps across supported platforms can reduce repeated data preparation and help researchers pursue focused questions while retaining the broader tissue context.

SpatialData provides a foundation for connecting these analyses by organizing expression measurements, images and other spatial elements within a common data framework. By recording the relationships between these elements and their coordinate systems, it allows expression data to remain linked to tissue structure during data selection and processing [7]. This shared representation can support workflows that apply established analysis methods to data from different platforms. Building on SpatialData therefore offers a way to connect broad platform support with further analysis of selected populations and regions.

Here, we present Spatialsnake, a flexible workflow that builds on SpatialData to support spatial transcriptomics analysis across sequencing- and imaging-based platforms (Fig. 1). Spatialsnake accepts processed outputs from Visium, Visium HD, Stereo-seq, Xenium and MERFISH and uses Snakemake to coordinate analysis steps through a common command-line interface [8]. Within this workflow, users can perform single- or multi-sample analysis, adjust preprocessing and clustering parameters, and select cell populations or tissue regions for further investigation.

**Figure 1.**
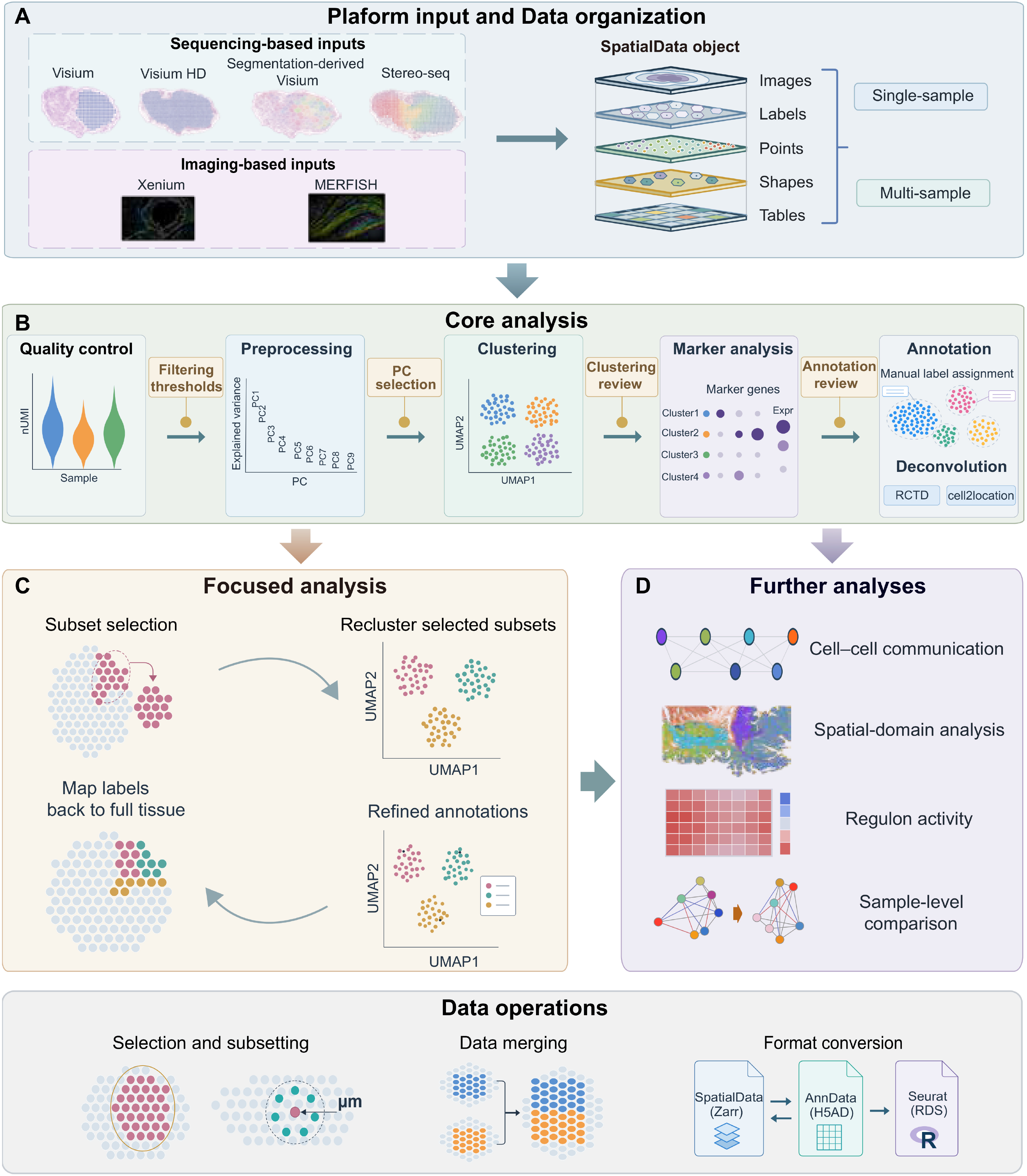
Overview of the Spatialsnake workflow. **(A)** Processed outputs from sequencing- and imaging-based spatial transcriptomics platforms are organized in SpatialData objects containing expression tables and available spatial elements, including images, labels, points and shapes. Single- and multi-sample datasets provide inputs to the analysis workflow. **(B)** Core analysis connects quality control, preprocessing, clustering, marker analysis, annotation and cell-type deconvolution. Users can adjust parameters and inspect saved intermediate results, while Snakemake manages dependencies and selective rerunning. **(C)** Populations, tissue regions or distance-based neighborhoods can be selected for further analysis. Selected subsets can be reclustered and annotated, and refined labels can be mapped back to the full dataset. **(D)** Additional modules support cell-type composition estimation, spatial-domain identification, cell–cell communication inference, regulon analysis and sample-level comparisons, using complete datasets or selected subsets as appropriate for each method. The bottom panel shows optional data operations, including selection and subsetting, merging, and conversion between supported SpatialData, AnnData and Seurat formats. Plots and tissue representations are schematic.

Selected subsets can be reclustered and annotated, and the refined labels can be mapped back to the complete tissue. The resulting datasets can then be used for cell-type composition estimation, spatial-domain identification, cell–cell communication inference, regulon analysis and sample-level comparisons, according to the input requirements of each method. Saved intermediate objects allow users to revise selected steps and extend their analyses without repeating the entire workflow. We demonstrate these capabilities using public spatial transcriptomics datasets, including a joint analysis of three colorectal cancer Visium HD sections that connects tissue-wide annotation with epithelial refinement. These examples illustrate how Spatialsnake supports both routine analysis and focused investigation of populations and regions within a common workflow.

### Fingdings

#### Spatialsnake provides a common workflow for spatial transcriptomics analysis across platforms

Spatialsnake brings processed spatial transcriptomics data from sequencing- and imaging-based platforms into a common analysis workflow (Fig. 1). It supports Visium, Visium HD, segmentation-derived Visium, Stereo-seq, Xenium and MERFISH, accommodating expression measurements associated with spots, bins or segmented cells. During data loading, expression measurements and sample metadata are organized with the available images, coordinates and other spatial elements in SpatialData objects. These objects provide a common starting point for single- and multi-sample analysis while retaining the spatial information supplied by each dataset. From this starting point, users can perform quality control, preprocessing, clustering and annotation through a common command-line interface. Analysis settings can be adjusted as results are inspected, and intermediate objects are saved for further use.

Users can then select an annotated population, a tissue region or observations within a specified distance of a target population for closer investigation. Selected subsets can be reanalysed, and refined annotations can be mapped back to the full dataset. Additional modules connect these datasets to downstream methods, while utility tools support splitting, merging and conversion between supported formats. Snakemake manages dependencies between analysis steps and supports selective rerunning, allowing users to revise or extend an analysis from saved results.

#### Spatialsnake supports core analysis of single- and multi-sample datasets

Spatialsnake connects quality control, preprocessing, clustering and annotation through saved analysis objects, allowing users to inspect results and adjust settings as the analysis proceeds. We illustrate single-sample analysis using P2CRC, one of three colorectal cancer sections profiled with Visium HD by Oliveira et al.[9], at 8-μm bin resolution (Fig. 2A–E). The distribution of total counts provides an initial view of expression depth and helps users choose filtering thresholds (Fig. 2A). After preprocessing, the PCA explained-variance plot supports selection of the number of components used for neighborhood construction and clustering (Fig. 2B). The same UMAP embedding is then colored by total counts, detected-gene counts and cluster assignment, allowing users to examine the relationship between expression depth and cluster structure (Fig. 2C). Marker expression and functional enrichment help interpret the clusters, with GO enrichment results for cluster 2 shown as an example (Fig. 2D). The resulting annotations can be viewed over the tissue image to assess their spatial distribution and relationship to tissue structure (Fig. 2E).

**Figure 2.**
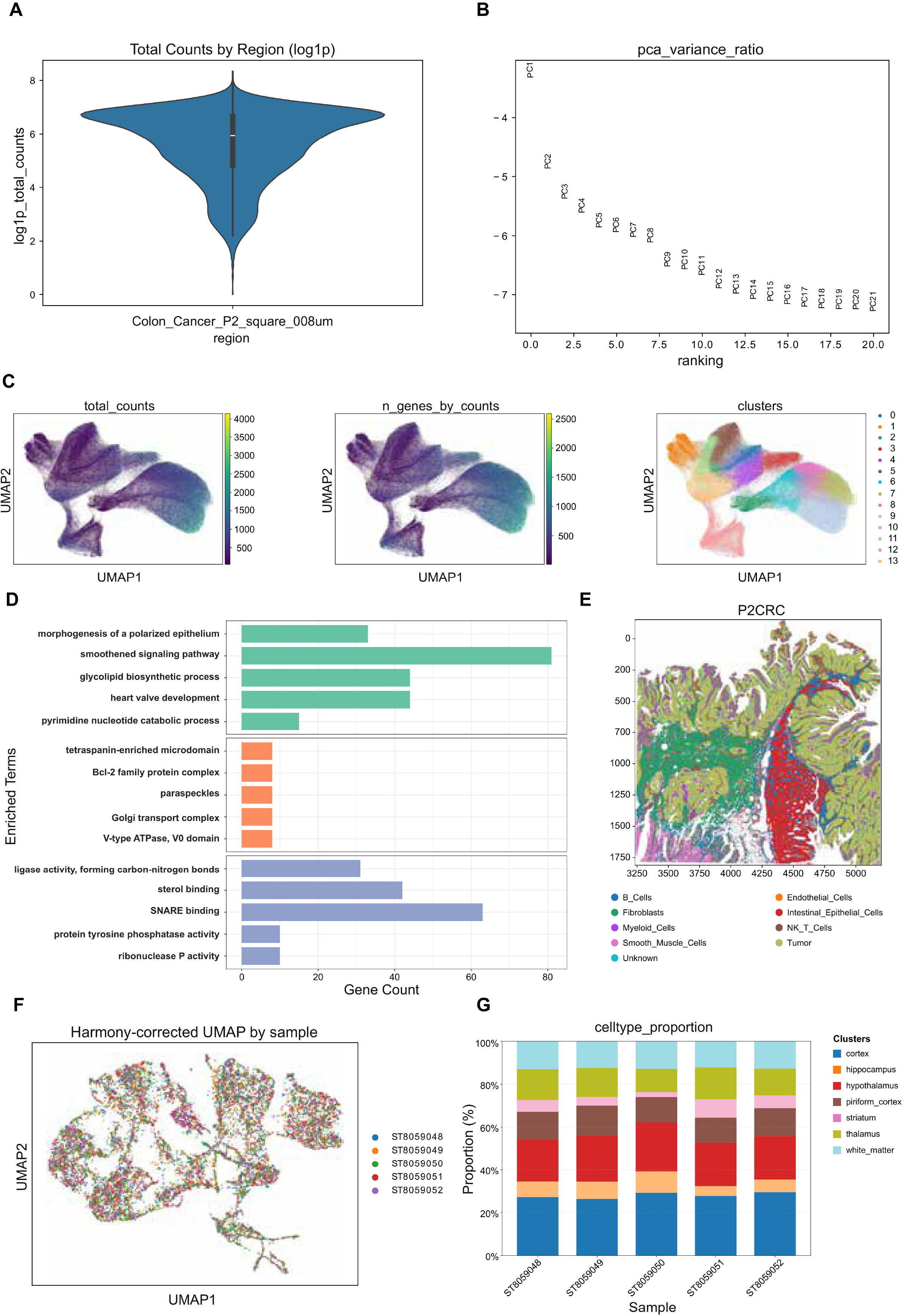
Representative outputs from single- and multi-sample core analyses. **(A)** Distribution of log1p-transformed total counts before filtering in CRC section P2CRC, analysed at 8-μm bin resolution. **(B)** Log-transformed PCA explained-variance ratios across the displayed components. **(C)** The same UMAP embedding colored by total counts, detected-gene counts and cluster assignment. **(D)** GO enrichment results for cluster 2, showing five terms from each of the biological process, cellular component and molecular function categories. Bar lengths indicate the number of genes associated with each term. **(E)** Reviewed spatial annotations overlaid on the tissue image. **(F)** Joint UMAP of five mouse-brain Visium sections, calculated using Harmony-adjusted principal components and colored by sample. **(G)** Proportions of spots assigned to each annotated brain region within each section.

For multi-sample analysis, Spatialsnake retains sample identities alongside the joint analysis results, allowing users to examine both the combined dataset and individual sections. We demonstrate this using five mouse-brain Visium sections, ST8059048– ST8059052, reported by Kleshchevnikov et al. [10] and available under ArrayExpress accession E-MTAB-11114. A joint UMAP calculated from Harmony-adjusted principal components shows the distribution of the five samples in a shared embedding (Fig. 2F). The annotations can then be summarized by section: Figure 2G shows the proportion of spots assigned to each annotated brain region. Together, these outputs allow researchers to inspect sample distributions after integration and examine tissue composition within each section using the same analysis object.

#### Spatialsnake supports focused analysis of populations and spatial neighborhoods

After examining tissue-wide annotations, users can select populations or regions for more foucsed analysis. Spatialsnake supports selection by sample, cluster, annotation or region of interest (ROI), as well as distance-based selection around a target population. For distance-based selection, users specify the targets, the distance range and, optionally, the annotations of eligible neighbors. Distances are calculated within each sample from each eligible observation to its nearest target, allowing users to examine either the broader surroundings of a population or its proximity to particular annotated groups.

We illustrate these options using section P2CRC from the CRC Visium HD dataset. Selecting tumor-labeled bins as targets and retaining all eligible non-target bins within 50 μm captures their surrounding tissue (Fig. 3A). Keeping the same targets and distance threshold while restricting neighbors to myeloid-labeled bins produces a more focused selection (Fig. 3B). Users can also define targets through a barcode list, as illustrated by a tumor-rich ROI and its neighbors within 100 μm (Fig. 3C). These distances are measured between bin locations, rather than from a geometric tumor boundary. The selected subsets retain their expression data, annotations and spatial coordinates for further analysis. Users can then repeat preprocessing, clustering and annotation as needed, and map refined labels back to the full dataset to examine them in their original tissue context.

**Figure 3.**
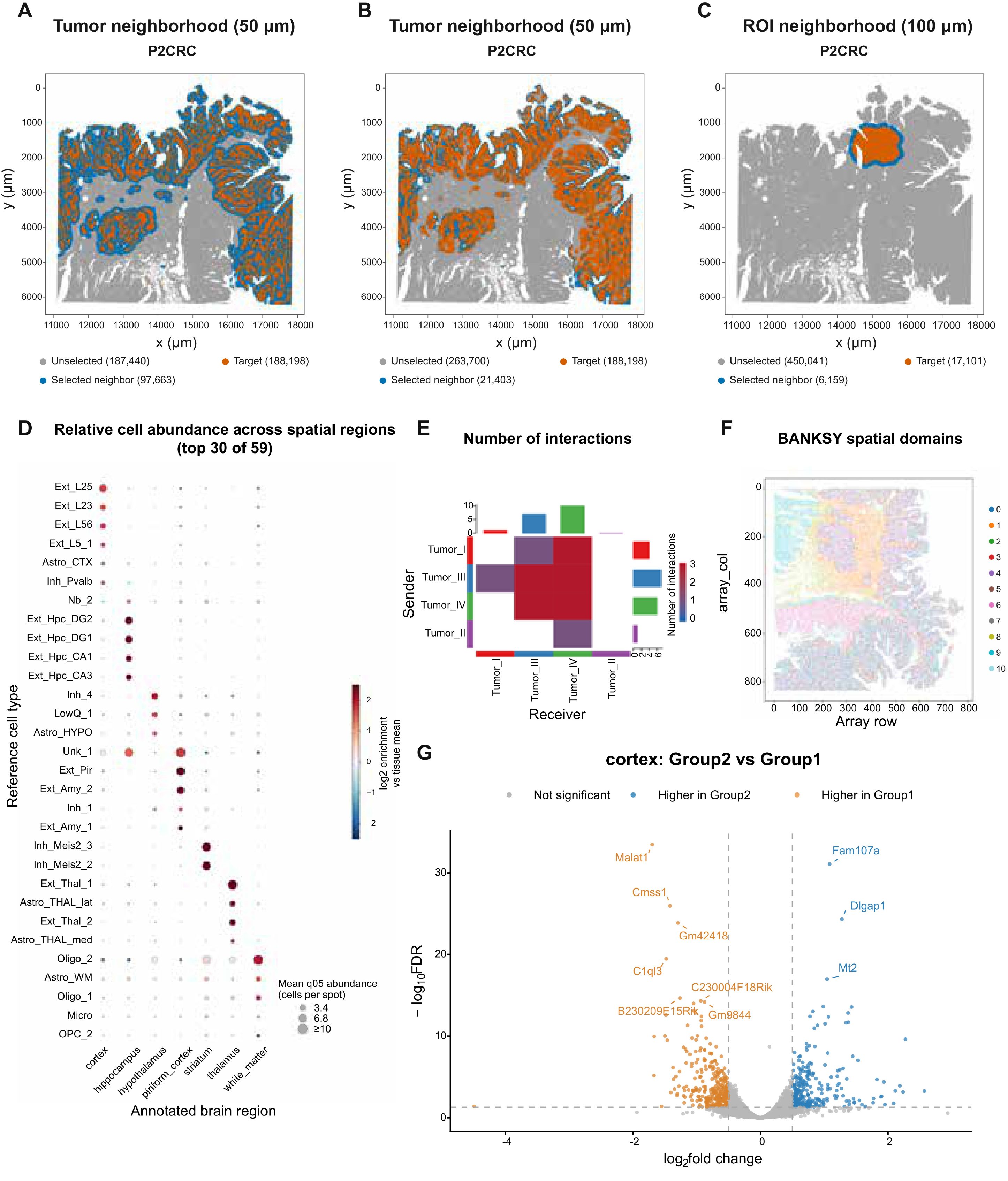
Focused spatial selection and downstream analyses with Spatialsnake. **(A)** Tumor-labeled target bins and non-target neighbors within 50 μm of their nearest target. **(B)** The same targets and distance threshold, with neighbors restricted to myeloid-labeled bins. **(C)** A barcode-defined tumor-rich region of interest and its non-target neighbors within 100 μm. In A–C, targets are orange, selected neighbors blue and unselected bins gray. Distances are measured between bin locations rather than from a geometric tumor boundary; coordinates are in micrometers. **(D)** Cell2location abundance estimates across seven annotated brain regions, showing the 30 reference cell types with the highest sample-balanced tissue mean abundance. Dot size represents the regional mean of posterior 5th-percentile abundance estimates, and color indicates log₂ enrichment relative to the sample-balanced tissue mean, clipped at ±2.5. **(E)** Spatial CellChat interaction counts among Tumor I–IV groups in P2CRC. Rows indicate senders and columns receivers; marginal bars show row and column totals. **(F)** BANKSY spatial-domain assignments in P2CRC, with 200,000 bins sampled for display using a fixed random seed. Axes show array-row and array-column coordinates. **(G)** Sample-level pseudobulk differential expression analysis of cortex spots, comparing Group2 with Group1. Group1 comprises ST8059048– ST8059050, and Group2 comprises ST8059051–ST8059052. Dashed lines indicate FDR = 0.05 and log₂ fold-change thresholds of ±0.5. Blue and orange indicate genes meeting both thresholds with higher expression in Group2 and Group1, respectively. All five sections share the same experimental condition; the grouping illustrates the comparison workflow.

#### Spatialsnake supports further analyses of annotated spatial datasets

Once spatial datasets have been annotated, researchers can investigate how the identified populations are distributed and how they relate to one another. Spatialsnake supports these investigations by preparing method-specific inputs from saved analysis objects. To examine cell-type composition, users can apply cell2location or RCTD with a compatible single-cell reference [11]. We demonstrate this using the five mouse-brain Visium sections, summarizing cell2location abundance estimates across seven annotated brain regions (Fig. 3D). The resulting profiles show how reference cell types are distributed across these regions.

Beyond population distributions, users can investigate potential communication between annotated groups using CellChat, CellPhoneDB or LIANA+[12–14]. In P2CRC, spatial CellChat summarizes candidate interactions among four tumor groups (Fig. 3E). Users can examine selected ligand–receptor pairs by replotting saved inference results, allowing different interactions to be explored without repeating inference (Supplementary Fig. S2). To examine tissue organization more broadly, BANKSY and CellCharter identify spatial domains using expression and spatial information[15,16], with BANKSY assignments in P2CRC shown in Figure 3F. At the level of gene regulation, pySCENIC provides gene regulatory network inference and regulon activity scoring [17], illustrated for endothelial and smooth-muscle groups in Supplementary Figure S1C.

These analyses characterize populations and their spatial relationships, while sample-level comparisons allow researchers to examine expression differences across groups of samples. Spatialsnake prepares these comparisons by aggregating counts within selected annotations for each sample. We illustrate the resulting pseudobulk analysis using cortex spots from the five mouse-brain sections, divided into groups of three and two samples (Fig. 3G). Because all five sections share the same experimental condition, this grouping demonstrates the comparison workflow rather than a treatment effect. Users can thus extend saved annotations into analyses of tissue composition, spatial organization, molecular interactions and sample-level expression differences through a common interface.

#### Spatialsnake connects tissue-wide annotation with epithelial subpopulation analysis

To demonstrate how core analysis and focused analysis can be used together, we jointly reanalysed the three CRC Visium HD sections, P1CRC, P2CRC and P5CRC, from Oliveira et al.[9]. Analysis of the 8-μm binned data retained 1,527,987 bins across the three sections. Broad annotations based on marker expression and tissue structure identified epithelial, tumor, stromal and immune groups, which were displayed separately for each section (Fig. 4A–C). PIGR, CEACAM6 and COL1A1 expression supported the intestinal epithelial, tumor and fibroblast annotations, respectively (Supplementary Fig. S1A), consistent with the source study.

**Figure 4.**
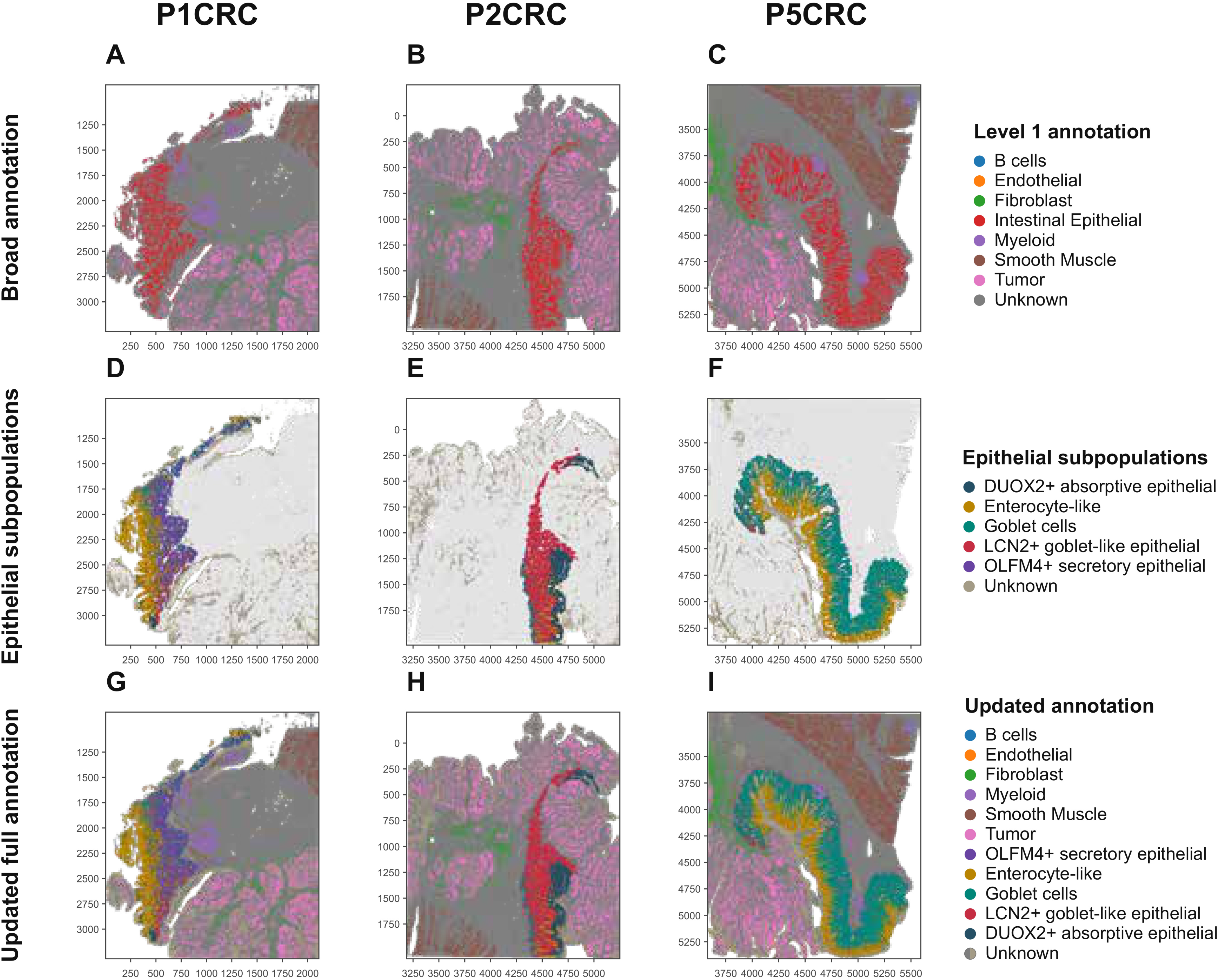
Tissue-wide annotation and epithelial subpopulation analysis in three colorectal cancer sections. Columns show P1CRC, P2CRC and P5CRC, analysed at 8-μm bin resolution. **(A–C)** Broad annotations based on marker expression and tissue structure, shown separately for each section. The combined dataset contains 1,527,987 retained bins. **(D–F)** Epithelial annotations after selection and reclustering of 223,632 intestinal epithelial bins into 13 subclusters. Marker expression supported five labels: OLFM4+ secretory epithelial, enterocyte-like, goblet, LCN2+ goblet-like epithelial and DUOX2+ absorptive epithelial. Subclusters without a clear assignment remain Unknown. **(G–I)** Refined epithelial labels mapped back to the full dataset, with non-target bins retaining their broad annotations. Annotations refer to spatial bins rather than individually segmented cells.

The intestinal epithelial group was then selected to examine differences that were not captured by the broad annotation. Reclustering its 223,632 bins produced 13 subclusters, whose marker expression supported five labels: OLFM4+ secretory epithelial, enterocyte-like, goblet, LCN2+ goblet-like epithelial and DUOX2+ absorptive epithelial. Subclusters without a clear assignment remained Unknown (Supplementary Fig. S1B). Goblet-associated groups expressed CLCA1, FCGBP and MUC2, while the LCN2+ goblet-like group also expressed REG1A and REG1B. Mapping these labels to their tissue locations showed different distributions across the sections: OLFM4+ secretory epithelial and enterocyte-like bins were prominent in P1CRC, LCN2+ goblet-like bins in P2CRC, and goblet-labeled bins in P5CRC (Fig. 4D–F).

To examine these epithelial groups alongside the rest of the tissue, we mapped their labels back to the full dataset while retaining the broad annotations of non-target bins (Fig. 4G–I). The updated maps show the epithelial subgroups together with surrounding tumor, stromal and immune groups. This example demonstrates how Spatialsnake connects joint analysis of complete sections with reclustering and annotation of a selected population, while keeping the results linked to their original tissue locations. The annotations describe groups of 8-μm bins rather than individually segmented cells.

#### Comparison with representative spatial transcriptomics workflows

We compared Spatialsnake with Spacemake, Panpipes and Sopa across platform support, core analysis, downstream methods and data operations (Table 1). The workflows share several functions, including quality control, annotation support and visualization, but differ in the technologies they support and the analyses available within each workflow. These differences affect how much additional data preparation and scripting users need when extending an analysis.

**Table 1.**
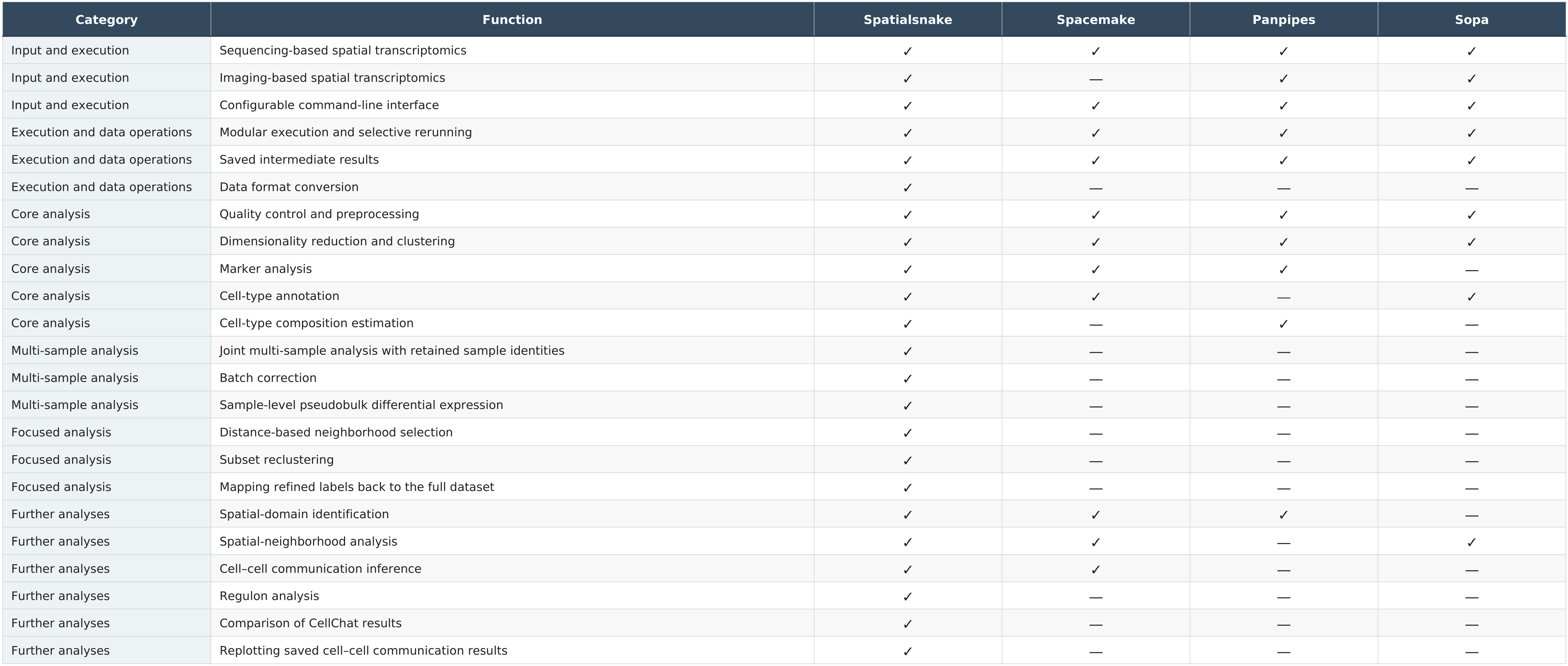
Comparison of supported functions across spatial transcriptomics workflows. ✓ indicates documented support in the examined release; — indicates that support was not identified in the reviewed sources. The comparison covers functions available through documented workflow commands or configuration options, including integrated third-party methods. Sources include the Spacemake, Panpipes and Sopa publications and documentation for the examined releases. The table compares functional coverage rather than analytical accuracy or computational performance.

Spatialsnake combines support for sequencing- and imaging-based inputs with adjustable analysis settings and further analysis of selected populations and regions. Users can select subsets, repeat clustering and annotation, map refined labels back to the full dataset, and prepare inputs for downstream methods through the same interface. Saved analysis objects connect these steps, while merging and format conversion support further use with other tools. This combination is useful for studies that move from tissue-wide annotation to the analysis of particular populations or spatial neighborhoods.

## Methods

### Implementation and execution

Spatialsnake can be installed using pip from PyPI or Conda from Bioconda. Its command-line interface is written in Python, and its main workflow is implemented in Snakemake. Users select modules such as Ingestion, Preprocess, Clustering and Annotation through the command-line interface. YAML files specify analysis methods, parameters and computing resources, with explicitly supplied command-line arguments taking precedence. Snakemake manages task dependencies and resources and calls Python and R scripts. Saved intermediate objects support parameter revision and selective rerunning. Downstream modules such as cell–cell communication analysis separate inference from plotting, allowing saved results to be reused when adjusting the displayed populations, pathways or ligand–receptor interactions.

Configuration records and execution logs document settings and run status. Data loading and organization (Ingestion) The Ingestion module imports processed platform outputs, including Space Ranger results for Visium, Visium HD and segmentation-derived Visium, SAW results for Stereo-seq, and outputs from the corresponding software for Xenium and MERSCOPE/MERFISH. Users retain the supported output directory structure and supply input paths, sample identifiers and required group assignments. Bin resolution or segmentation-derived input is selected before loading. Available images, transcript positions and cell boundaries are imported according to reader settings.

SpatialData objects store expression matrices and observation annotations in AnnData tables, alongside spatial elements representing images, segmentation labels, transcript positions and cell or capture-region geometries. Identifiers link tables to the spatial elements they annotate, while coordinate transformations relate the spatial elements. Zarr storage supports reuse across modules. Multi-sample loading combines expression tables while retaining sample and group identities and distinct spatial-element names. Concatenation and optional batch correction are performed separately.

Ingestion also calculates quality-control (QC) metrics and generates diagnostic plots of total counts, detected genes and mitochondrial counts, together with the relationship between counts and detected genes. These summaries allow users to inspect distributions within and across samples and choose filtering thresholds for preprocessing.

### Datasets

Two published datasets were used. The colorectal cancer (CRC) Visium HD dataset comprised P1CRC, P2CRC and P5CRC at 8-μm bin resolution. P2CRC supported the single-sample and selected downstream analyses in Figures 2 and 3; all three sections supported the joint analysis and epithelial refinement in Figure 4. Five mouse-brain Visium sections, ST8059048–ST8059052, from ArrayExpress E-MTAB-11114 supported multi-sample and selected downstream analyses in Figures 2 and 3. Download sources are provided in the Data availability statement.

### Preprocessing, clustering and annotation

Preprocessing and clustering use Scanpy [18]. Observations represent spots, bins or segmented cells. QC filters specify minimum total counts per observation, maximum mitochondrial count percentages and the minimum number of observations detecting each gene. Samples can be filtered separately, and the intersection of retained genes is used for joint analysis. Raw counts are preserved. Expression is normalized to 10,000 counts per observation and log1p-transformed, with optional highly variable gene selection before principal component analysis (PCA).

The PCA explained-variance plot guides component selection. A separate heuristic expresses each component’s standard deviation as a percentage of the sum across components. It recommends the smaller of two cutoffs: the first component with cumulative contribution above 90% and individual contribution below 5%, or the component following the last adjacent decrease above 0.1 percentage points. Users specify clustering dimensionality after reviewing the plot and logged recommendation.

Multi-sample analyses optionally use Harmony-adjusted PCA coordinates[19] or a BBKNN batch-aware neighborhood graph[20]. For large datasets, GeoSketch can select reference subsets within samples using PCA or Harmony coordinates[21]. Leiden[22] and Louvain[23] cluster the neighborhood graph, whereas k-means uses the selected low-dimensional representation; UMAP[24] provides visualization. In sketch mode, Scanpy ingest maps reference cluster labels and UMAP coordinates to the full dataset, with checks of gene alignment, representations, observation counts and label completeness.

Scanpy differential expression testing identifies candidate marker genes. Marker expression, GO/KEGG enrichment[25] and spatial plots guide manual annotation. Users assign cell identities to cluster identifiers in an annotation table. The workflow maps these identities to the corresponding observations, saves the annotated object and generates spatial plots, UMAP plots and plots of annotation proportions. These outputs guide further annotation or parameter revision.

For mixed locations, cell2location and RCTD estimate cell-type composition from annotated single-cell reference counts with compatible gene identifiers [10,11]. cell2location uses reference expression signatures and a Bayesian model to estimate abundances, with expected cells per location and detection-efficiency variation priors set for the tissue and measurement scale. RCTD accounts for platform effects; full mode permits multiple cell types per location, whereas doublet mode permits at most two. Mode selection follows resolution and expected mixing, accommodating conventional Visium spots and Visium HD bins at the selected granularity.

Spatial maps and regional bubble plots summarize cell2location posterior 5th-percentile abundances, RCTD full-mode weights normalized per location or doublet-mode primary-label frequencies among classified locations. Multi-sample summaries are calculated within samples and averaged equally across samples containing each region. Figure 3D displays the 30 reference types with the highest sample-balanced tissue-wide mean abundance across seven annotated brain regions.

### Spatial selection and focused analysis

Subsets can be selected by sample, group, cluster, cell-type annotation, observation identifiers or region of interest (ROI). These selection methods define inputs for focused analyses. To investigate local cellular environments, users can additionally select neighbors within specified distances of a target population or ROI and restrict candidate neighbors by annotation. Targets and selected neighbors remain linked to the original tissue for subsequent reclustering and annotation refinement.

Before distance-based selection, coordinates are converted to micrometers using sample-specific calibration, giving thresholds a consistent physical meaning across platforms and resolutions. Visium, Visium HD and segmentation-derived Visium use full-resolution image coordinates multiplied by the physical length per pixel (microns_per_pixel). For conventional Visium lacking this value, calibration is approximated by the nominal 55-μm spot diameter divided by its recorded pixel diameter. Visium HD bin side length, such as 8 μm, defines measurement resolution; image coordinates still require pixel calibration. Segmentation-derived Visium uses cell-geometry centroids in the calibrated image coordinate system.

Stereo-seq calibration uses the DNA nanoball (DNB) spacing recorded by the GEF resolution parameter. Dividing this value in nanometers by 1,000 gives micrometers per native coordinate unit. For example, a 500-nm spacing corresponds to 0.5 μm per unit and a nominal bin50 side length of 25 μm. Regular-bin and cellbin coordinates use the spacing recorded in their input. Supported Xenium and MERSCOPE/MERFISH cell-center coordinates are already in micrometers, so their conversion factor is 1. Missing calibration, invalid coordinates or identifier mismatches stop selection.

For candidate i and target set Tₖ in the same sample, nearest-target distance is:

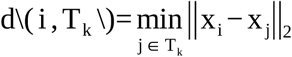

Here, xᵢ and xⱼ are two-dimensional micrometer coordinates. A cKDTree retains non-target candidates at or above the lower distance bound and below the upper bound, together with all targets. Distances are between observation locations or centroids within each sample, not from boundaries. ROI targets are identified by observation IDs or full coverage of linked geometries. Distance-based ROIs support Visium, Visium HD, segmentation-derived Visium and Xenium, one physical sample per run; polygon-coordinate ROIs in this mode are limited to Xenium.

The selected subset retains identifiers, sample metadata, annotations and spatial information and can directly enter reclustering. This step reuses preprocessed expression, optionally reselects highly variable genes, and performs PCA, neighborhood construction, UMAP and Leiden clustering with marker output. Revised quality-control or normalization settings require preprocessing first. Following marker and spatial review, refined labels are mapped back by observation and sample or region identifiers, with duplicate and conflict checks. Unselected observations retain their annotations. This links distance-based selection, subgroup refinement and annotation merging within the original tissue context.

### Downstream analyses

Downstream modules analyze user-selected full tissues, regions or refined populations. Spatial-domain identification uses BANKSY through pyBANKSY, which combines expression and local neighborhood features, or CellCharter, which aggregates latent expression features over spatial neighborhoods [15,16]. Multi-sample analyses construct spatial neighborhoods within each sample. Figure 3F displays 200,000 bins sampled with a fixed random seed.

Transcription factor network inference and regulon activity analysis use pySCENIC [17], with GRNBoost2 for candidate co-expression relationships, motif enrichment for regulon definition and AUCell for activity scoring. Users provide species-compatible transcription factor lists, motif-ranking databases and motif annotations. Cell–cell communication inference uses CellChat, CellPhoneDB or LIANA+ with expression data, population labels and compatible ligand–receptor resources [12–14].

For spatial CellChat [12], Spatialsnake matches sample-specific coordinate conversion factors and effective observation sizes to each input. Visium platforms use image pixel calibration, Stereo-seq uses DNB spacing, and micrometer-based Xenium and MERSCOPE/MERFISH coordinates use a factor of 1.

The effective size is the nominal 55-μm spot diameter for conventional Visium, the selected bin side length for Visium HD, or grid side length multiplied by DNB spacing in micrometers for regular-bin Stereo-seq. Segmentation-derived Visium, Stereo-seq cellbin, Xenium and MERSCOPE/MERFISH use a configurable size defaulting to 10 μm. Distance tolerance is half this size. Native coordinates and their conversion factors are passed together to CellChat. The spatial interaction range defaults to 250 μm and can be adjusted for the tissue and research question. Inferred interactions, networks and pathways are saved for subsequent visualization of selected populations and signals.

### Sample-level comparisons

Raw counts for each selected cell type or region are summed by gene within samples to form pseudobulk matrices. Counts must be non-negative integers; conditions come from the sample table. Samples are filtered by target-observation counts, genes by total counts and the number of samples with nonzero counts. At least two samples per condition must remain or analysis stops. Samples serve as the units of analysis.

PyDESeq2[26] or edgeR[27] tests condition contrasts with condition as the only design factor. PyDESeq2 fits DESeq2 count models; edgeR applies expression filtering, TMM normalization, dispersion estimation and quasi-likelihood F-tests. Outputs include log₂ fold changes, P values, adjusted P values, volcano plots, heatmaps and GO/KEGG enrichment.

Figure 3G aggregates cortex spots per mouse-brain section, comparing Group1 (ST8059048–ST8059050) with Group2 (ST8059051–ST8059052) at FDR < 0.05 and absolute log₂ fold change > 0.5. This workflow demonstration groups sections from the same experimental condition.

CellChat comparisons use separately fitted objects with compatible databases, data types and population labels. The comparison module summarizes differences in interaction counts, interaction strengths, signaling pathways and ligand–receptor interactions between the fitted networks.

### Data merging and format conversion

Sample merging aligns expression matrices by gene intersection or union while retaining sample identifiers, annotations and spatial elements. Annotation merging instead maps refined labels using observation and sample or region identifiers without changing expression data or spatial structure. Both support reuse across modules.

Conversion supports SpatialData/Zarr export to H5AD or Seurat/RDS, and H5AD import into SpatialData/Zarr or conversion to Seurat/RDS. Selected, validated raw counts enter the Seurat counts layer; other expression values enter its data layer. schard transfers compatible Visium images and scales; other coordinates are stored in a Seurat dimensional-reduction object. RDS files are reloaded to check class and dimensions, with source matrices and routes recorded. H5AD import retains supplied spatial information or creates a table-only SpatialData object.

### Three-section CRC reanalysis and resource measurements

The joint CRC analysis used P1CRC, P2CRC and P5CRC at 8-μm bin resolution [9]. Following the preprocessing described above, Harmony adjusted the low-dimensional representation, and GeoSketch selected 30% of observations per section. The 17 reference clusters were mapped to the full dataset and broadly annotated using marker expression and tissue locations.

The 223,632 intestinal epithelial bins were reclustered into 13 subclusters. Marker inspection and manual review assigned interpretable groups to five epithelial categories, with unresolved groups labeled Unknown. Refined labels were mapped back by observation and sample or region identifiers, retaining broad annotations and checking the results in complete sections.

Runtime and peak resident set size (peak RSS) were recorded on a Linux server with an AMD EPYC 9654 processor and four configured CPU threads. Preprocessing took 58.3 min (26.2 GiB peak RSS), clustering 29.2 min (14.3 GiB), and epithelial reclustering 8.0 min (15.9 GiB). Clustering included reference clustering and full-dataset label mapping. These are single-run measurements under the stated settings.

## Conclusions

Spatialsnake brings data from sequencing- and imaging-based spatial transcriptomics platforms into a common analysis workflow. Built on SpatialData, it connects core analysis with population and region selection, reclustering, annotation mapping and downstream methods. Adjustable parameters and saved intermediate results allow researchers to revise and extend analyses as their questions develop.

Applications to mouse-brain Visium and colorectal cancer Visium HD data demonstrate both single- and multi-sample analysis. In particular, the joint analysis of three CRC sections shows how users can move from tissue-wide annotation to epithelial subpopulation analysis and map refined labels back to the complete tissue. These examples illustrate how Spatialsnake supports routine analysis and focused investigation within the same workflow.

Future development will expand support for spatial technologies and analysis methods and improve the handling of large, high-resolution datasets. Its modular design allows additional tools to be incorporated into the existing workflow.

## Authors’ contributions

SWX conceived and supervised the study. ZHL and ZHZ developed the software. ZHZ performed the data analyses. ZHL drafted the manuscript, and SWX reviewed and edited it. All authors read and approved the final manuscript.

## Funding

This work was supported by the Guangdong Province College Student Innovation Training Program (202610573002S and G202604302004) and the Guangdong Provincial Administration of Traditional Chinese Medicine TCM Research Projects (20261211).

## Data availability

The human colorectal cancer Visium HD data were obtained from the study by Oliveira et al. [9] and are available at https://www.10xgenomics.com/platforms/visium/product-family/dataset-human-crc. The mouse-brain Visium data were obtained from the study by Kleshchevnikov et al. [10] and are available from ArrayExpress under accession number E-MTAB-11114 at https://www.ebi.ac.uk/biostudies/arrayexpress/studies/E-MTAB-11114.

## Code availability

Spatialsnake source code is available at https://github.com/l-zh007/spatialsnake. Documentation and tutorials are available at https://spatialsnake-tutorial.readthedocs.io/en/latest/index.html. The package is distributed through PyPI at https://pypi.org/project/spatialsnake/ and Bioconda at https://anaconda.org/channels/bioconda/packages/spatialsnake/overview.

## Declarations

### Ethics approval and consent to participate

Not applicable. This study used publicly available datasets and involved no new recruitment of human participants or animal experiments.

## Consent for publication

Not applicable. Competing interests

The authors declare no competing interests.

## Notes

### Competing Interest Statement

The authors have declared no competing interest.

### Summary of Updates

The main text of the paper remains unchanged, with only the author name format corrected.

