## Supplementary Materials for "Spatialsnake: a flexible workflow for spatial transcriptomics analysis across platforms"

**Contents**

Supplementary Figure S1 (A–C). Marker-expression dot plots for broad and epithelial annotations, and a pySCENIC regulon-activity heatmap.

Supplementary Figure S2 (A–B). Selected ligand–receptor interactions replotted from the same saved LIANA inference results.

Supplementary Table S1. Core software versions and workflow modules.

### Supplementary Figure S1A | Broad cell-type marker expression

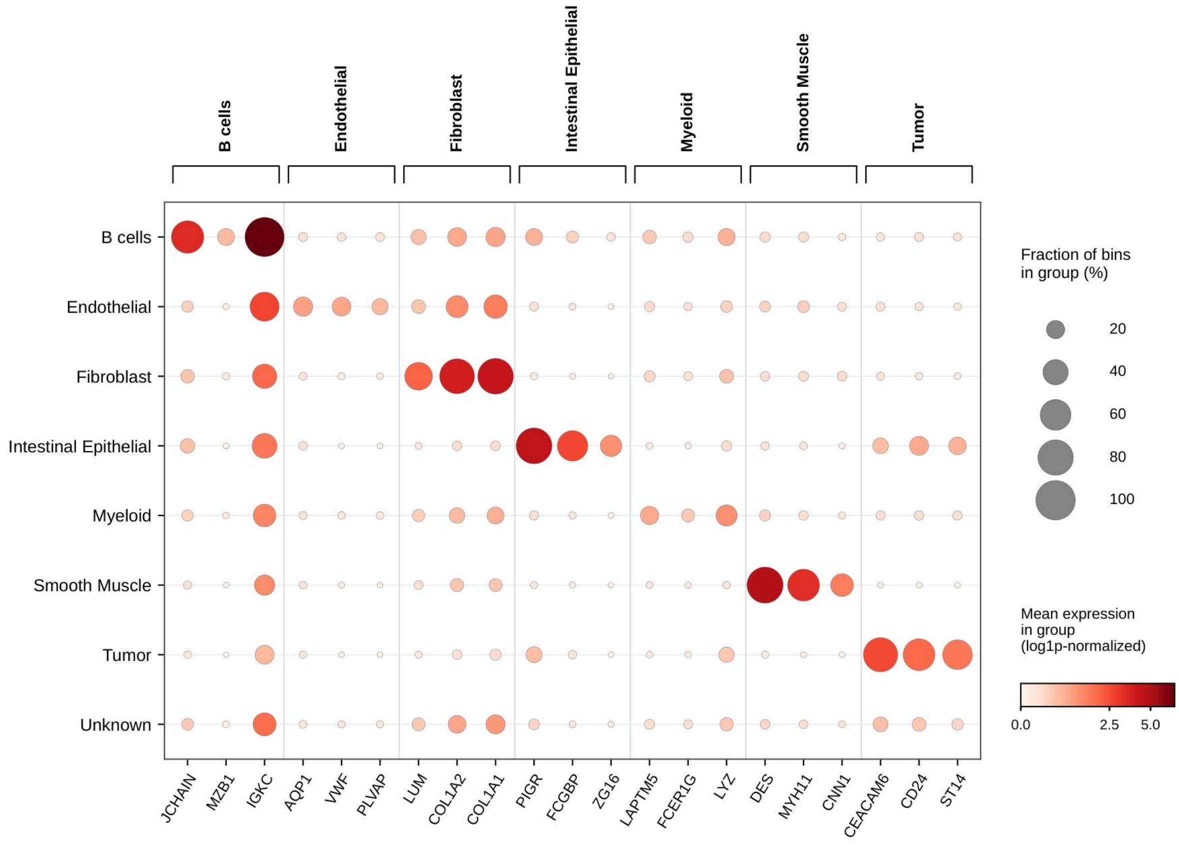


**Supplementary Figure S1A.** Representative marker-expression dot plot for the broad CRC Visium HD annotation classes B cells, Endothelial, Fibroblast, Intestinal Epithelial, Myeloid, Smooth Muscle, Tumor, and Unknown. Rows indicate the eight retained working annotation categories; columns are genes grouped under their corresponding marker categories. Dot size represents the percentage of spatial bins with detectable expression in each group, and dot colour denotes mean expression on the log1p-normalized scale shown in the source plot. The Unknown category is included for completeness; these patterns support review of expression-based annotations but do not establish one-to-one cell identities for Visium HD bins.

**Supplementary Figure S1B | Epithelial computational subcluster markers**

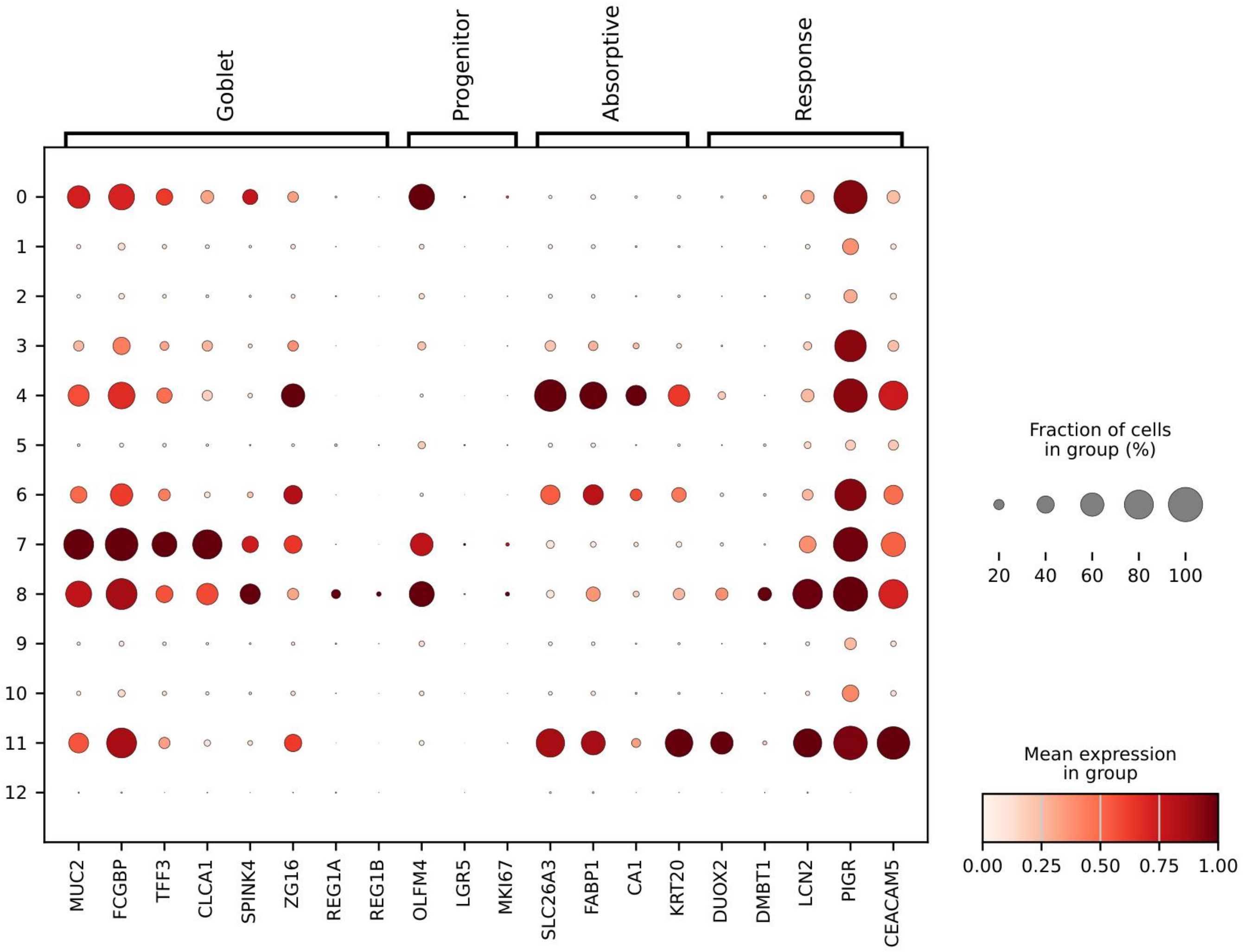


**Supplementary Figure S1B.** Marker-expression dot plot across 13 computational clusters (0–12) from the selected intestinal epithelial compartment in the CRC Visium HD reanalysis. Twenty representative genes are grouped as Goblet, Progenitor, Absorptive, and Response markers. The four previously included Lineage/QC genes (CXCL8, S100A8, PTPRC, and HBA2) are omitted from this updated view. Dot size follows the source plot's fraction-expressing percentage, and dot colour uses its mean-expression display scale (0–1). Although the source size legend uses the term "cells," the relevant analytical observations are spatial bins. These expression patterns support annotation review rather than independently confirm cell identity.

### Supplementary Figure S1C | pySCENIC regulon activity in endothelial and smooth-muscle groups

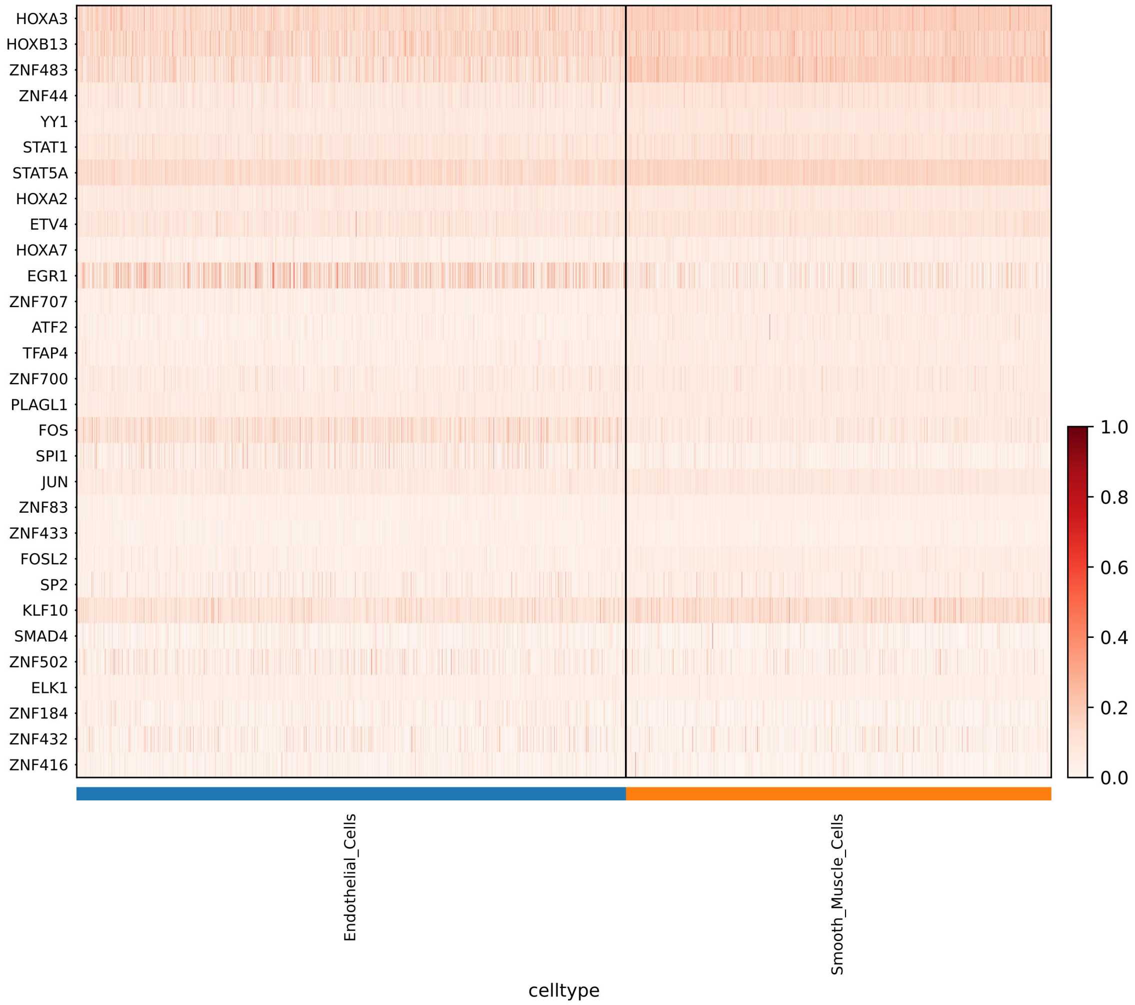


Supplementary Figure S1C. pySCENIC AUC heatmap for the displayed Endothelial_Cells and Smooth_Muscle_Cells annotation groups. Rows show the transcription-factor-associated regulons identified in the plot (including HOXA3, STAT1 and FOS), and columns show plotted observations arranged by the two group labels. The bar beneath the heatmap identifies the groups; the colour key displays a 0–1 scale. This panel illustrates a regulon-activity summary, not a marker-gene expression matrix. The analysis object, regulon-selection rule and any AUC rescaling should be checked against the original plotting record before submission.

### Supplementary Figure S2A | EREG/AREG interaction selection from saved LIANA results

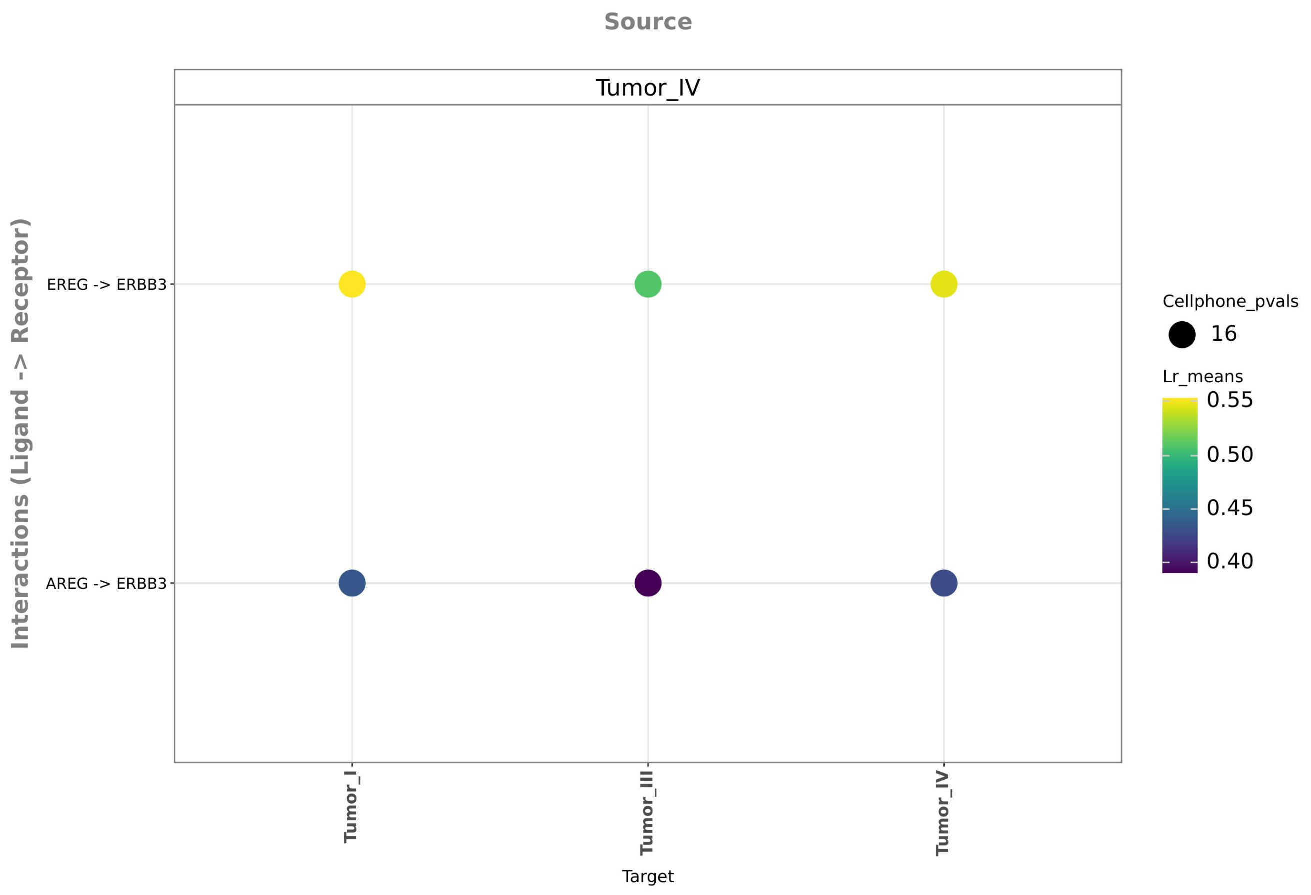


**Supplementary Figure S2A.** Focused ligand–receptor dot plot from the saved LIANA inference table for the CRC P2 working Tumor I–IV groups. Tumor IV is the sender and Tumor I–IV are candidate receivers. Selecting EREG–ERBB3 and AREG–ERBB3 using liana_pairs displays six sender–receiver–interaction records. Dot colour represents the stored lr_means score, while dot size represents the transformed CellPhoneDB permutation p-value used by the plotting function. The displayed saved p-values are zero, resulting in equal-sized dots. This view reuses the original 438-row inference table without rerunning LIANA.

### Supplementary Figure S2B | VEGFA interaction selection from saved LIANA results

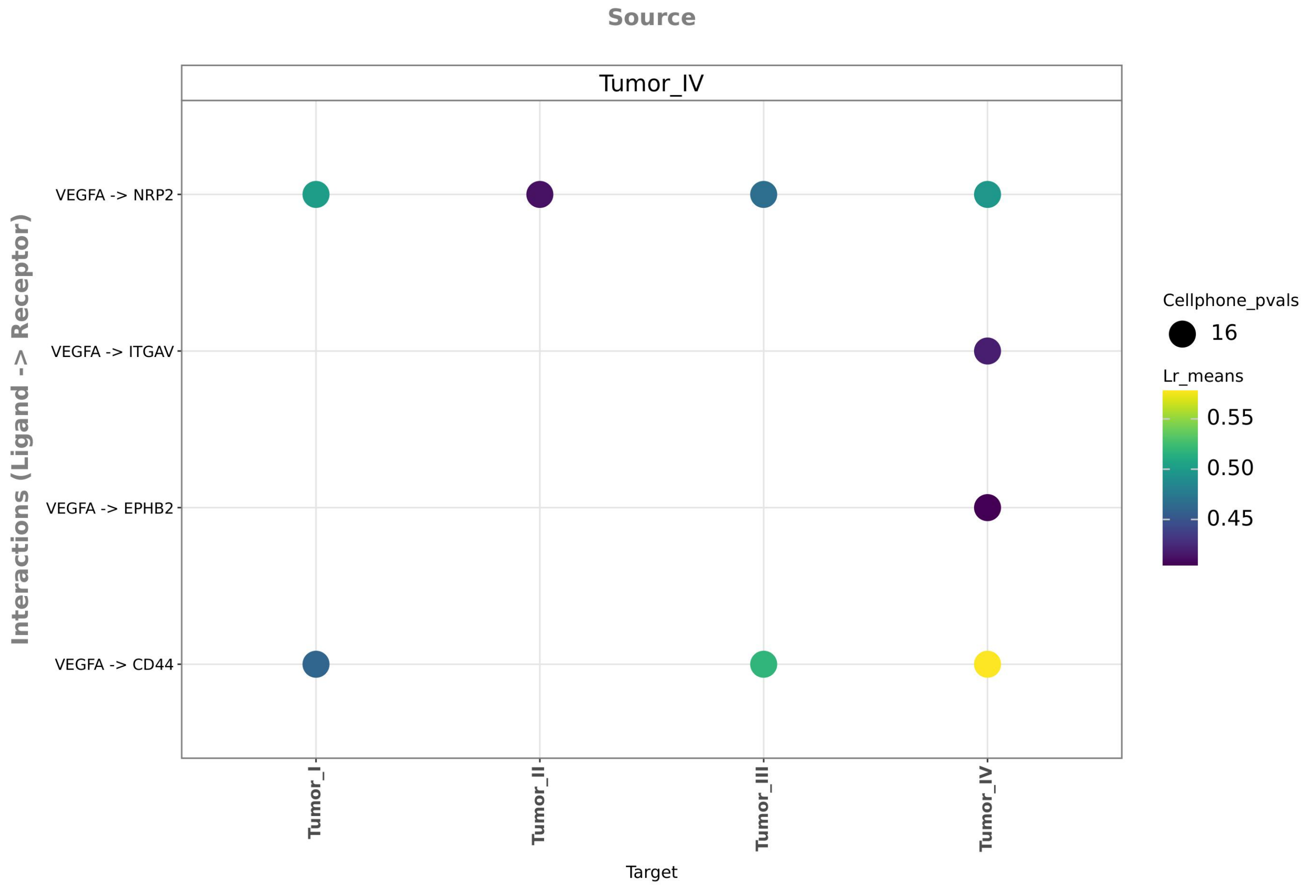


**Supplementary Figure S2B.** The same saved LIANA inference table, Tumor IV sender, Tumor I–IV receivers, liana_top_n of 6, and plotting p-value cutoff of 0.05 were used as in Supplementary Figure S2A. Only liana_pairs was changed to select VEGFA–NRP2, VEGFA–CD44, VEGFA–ITGAV, and VEGFA–EPHB2, yielding nine displayed records. Dot size and colour follow the encodings described for S2A. The native colour scale is determined independently for each view; colours should therefore not be interpreted as directly comparable interaction strengths across panels. Together, S2A and S2B illustrate focused replotting of previously inferred interactions.

### Supplementary Table S1 | Core software versions and workflow modules

| Software | Version | Workflow module |
|---|---|---|
| Snakemake | 9.8.1 | Workflow execution and dependency management |
| SpatialData / spatialdata-io | 0.5.0 / 0.3.0 | Platform ingestion and spatial data organization |
| Scanpy | 1.10.4 | Preprocessing, clustering, marker analysis, and label projection |
| Harmony (harmonypy) | 0.0.10 | Multi-sample integration |
| BBKNN | 1.6.0 | Batch-aware neighborhood construction |
| GeoSketch | 1.3 | Reference sampling |
| cell2location | 0.1.5 | Reference-assisted annotation |
| RCTD (spacexr) | 2.2.1 | Reference-assisted annotation |
| BANKSY | 1.3.4 | Spatial-domain analysis |
| CellCharter | 0.3.5 | Spatial-domain analysis |
| CellChat | 1.6.1 | Communication inference, visualization, and comparison |
| LIANA | 1.6.1 | Ligand–receptor inference and visualization |
| CellPhoneDB | 5.0.1 | Ligand–receptor inference |
| pySCENIC | 0.12.1+11.g06bafba | Regulon inference and activity scoring |
| PyDESeq2 | 0.5.2 | Sample-level pseudobulk gene comparison |
| edgeR | 4.4.2 | Sample-level pseudobulk gene comparison |
| clusterProfiler | 4.14.6 | Functional enrichment |
| schard / Seurat | 1.1.0 / 5.5.0 | Export and validation of Seurat objects |

**Table legend.** Versions were checked on 13 October 2026. Python 3.12.11 and R 4.4.2 were used in the examined demonstration environment. These version records do not describe the historical three-section CRC application (Spatialsnake 0.0.4 with local changes). Inclusion of a software package in this table does not imply that it was executed in every analysis example.
